# ColorBayes: Improved color correction of high-throughput plant phenotyping images to account for local illumination differences

**DOI:** 10.1101/2022.03.01.482532

**Authors:** Diego Lozano-Claros, Eddie Custovic, Guang Deng, James Whelan, Mathew G. Lewsey

## Abstract

**Background:** Color distortion is an inherent problem in image-based phenotyping systems that are illuminated by artificial light. This distortion is problematic when examining plants because it can cause data to be incorrectly interpreted. One of the leading causes of color distortion is the non-uniform spectral and spatial distribution of artificial light. However, color correction algorithms currently used in plant phenotyping assume that a single and uniform illuminant causes color distortion. These algorithms are consequently inadequate to correct the local color distortion caused by multiple illuminants common in plant phenotyping systems, such as fluorescent tubes and LED light arrays. We describe here a color constancy algorithm, ColorBayes, based on Bayesian inference that corrects local color distortions. The algorithm estimates the local illuminants using the Bayes’ rule, the maximum a posteriori, the observed image data, and prior illuminant information. The prior is obtained from light measurements and Macbeth ColorChecker charts located on the scene.

**Results:** The ColorBayes algorithm improved the accuracy of plant color on images taken by an indoor plant phenotyping system. Compared with existing approaches, it gave the most accurate metric results when correcting images from a dataset of Arabidopsis thaliana images.

The software is available at https://github.com/diloc/Color_correction.git.

## Background

Leaf color is a plant trait that can be used as an indicator for plant nutrition, water absorption and transpiration, photosynthetic efficiency, and performance [1-3]. This trait is linked with the amount and concentration of pigments within leaves, which determine their color and indicate leaf physiology [4]. For example, plants growing in nitrogen (N) deficiency have characteristic light-green and yellow leaves. In contrast, deep green color in stunted plants can suggest toxicity symptoms due to excessive N concentration [5].

There are numerous manual and automatic techniques to measure leaf color. The manual methods involve visual inspection and matching with a reference such as the Royal Horticultural Society Color chart and Munsell charts [6]. However, these approaches are destructive, time-consuming, and not cost-effective. Automatic and semi-automatic techniques can measure leaf color with minimal human intervention and overcome limitations posed by manual processes. For instance, aerial and satellite imagery with limited resolution is suitable for measuring the foliage color of large crop fields and forests [7]. Meanwhile, high throughput phenotyping systems can measure the individual leaf color of thousands of plants located in outdoor and indoor environments. However, varying illumination conditions can affect image-based methods by altering the measured color of pixels [8, 9]. This distorted image data is no longer a faithful representation of the true leaf color and may cause incorrect biological interpretation of the data [10].

Color constancy describes the ability to identify the color of objects irrespective of lighting conditions [11]. In computer vision, color constancy is challenging since algorithms must estimate the unknown illumination parameters and calculate their effect on the scene using surface descriptors [12, 13]. Some assumptions are usually made to simplify this complex problem, including a single invariant canonical illuminant, a balanced camera, and ideal diffuse reflecting surfaces [14]. Additional factors may be considered, like surface reflectance characteristics, camera response, and illuminant distribution.

Color constancy algorithms can be classified into the following categories: color calibration, grey world, gamut mapping, Retinex, probabilistic methods, neural networks, and automatic color equalization [15-22]. Color calibration algorithms are the most commonly used method to correct color distortion in visual range image-based plant phenotyping [8]. In this approach, images are corrected by adjusting the deviation between a color rendition chart and its color response on the recorded image. The chart is often referred to as Macbeth ColorChecker, a card that displays 24 reference patches of known, defined colors under a standard illuminant [23].

The above color constancy algorithms correct image colors by assuming a single and uniform canonical illuminant. However, this assumption cannot be made for many high-throughput phenotyping systems as they frequently use multiple artificial light sources that produce non-uniform light distribution. The uncorrected leaf color of plants in images from such systems will follow a similar pattern to the non-uniform illuminant distribution, which causes local color distortion and subsequent inaccurate reporting of plant color.

We present a model based on Bayesian inference that corrects the color distortion caused by non-canonical illuminants in plant image data. The impact of local color distortion was reduced by applying the image correction, and, as a result, the color of leaves was more accurately represented than with other color constancy methods commonly used in plant phenotyping.

## Results

The main objective of the study was to correctly identify plant color under the non-uniform artificial light of high throughput plant phenotyping (HTPP) systems. This was accomplished by developing a color correction algorithm based on Bayesian inference and a ground truth image dataset. Our Bayesian algorithm increased the accuracy of leaf color measurements from images compared to other color constancy algorithms commonly used to correct HTPP systems. We tested the algorithm on experiments comprised of *Arabidopsis thaliana* plants (ecotype Col-0). One hundred and forty-eight *A. thaliana* individuals were used in the experiment. An image-based HTPP system with fixed plant positions was set to take images of plants every 15 minutes from the 13^th^ to the 35^th^ day after sowing (DAS). The HTPP system consisted of multiple cameras, fixed plant positions, and multiple LED illumination sources [24]. The measured leaf color of individual plants varied across the imaging chamber (**Figure 1**). Variation in measured color was not consistent with the human observation of plant color. This inconsistency occurs because there is variation in local lighting conditions, a common limitation in fixed-position, artificially illuminated HTPP systems (**Figure 2**).

**Figure 1.**
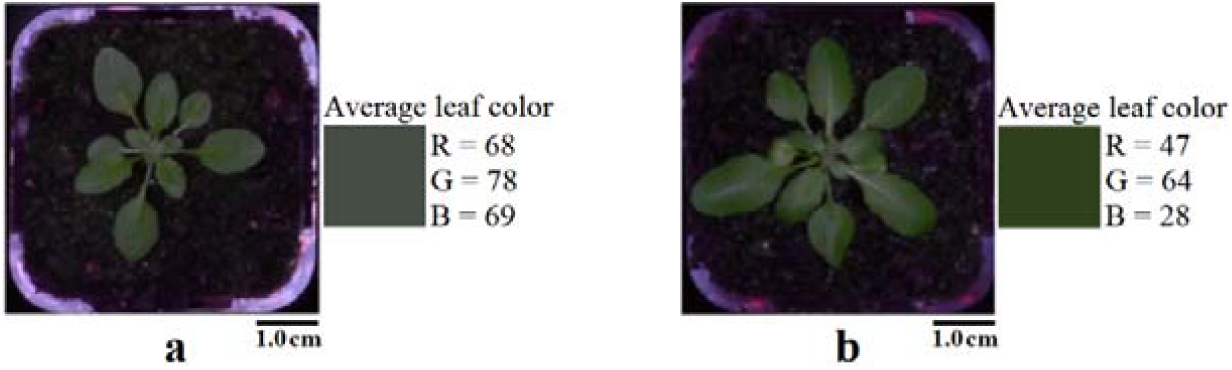
Leaf color difference caused by distinct local illuminants. (a) A plant with light leaf color appearance and (b) a plant with dark leaf color appearance, both imaged within the same experiment at the same time

**Figure 2.**
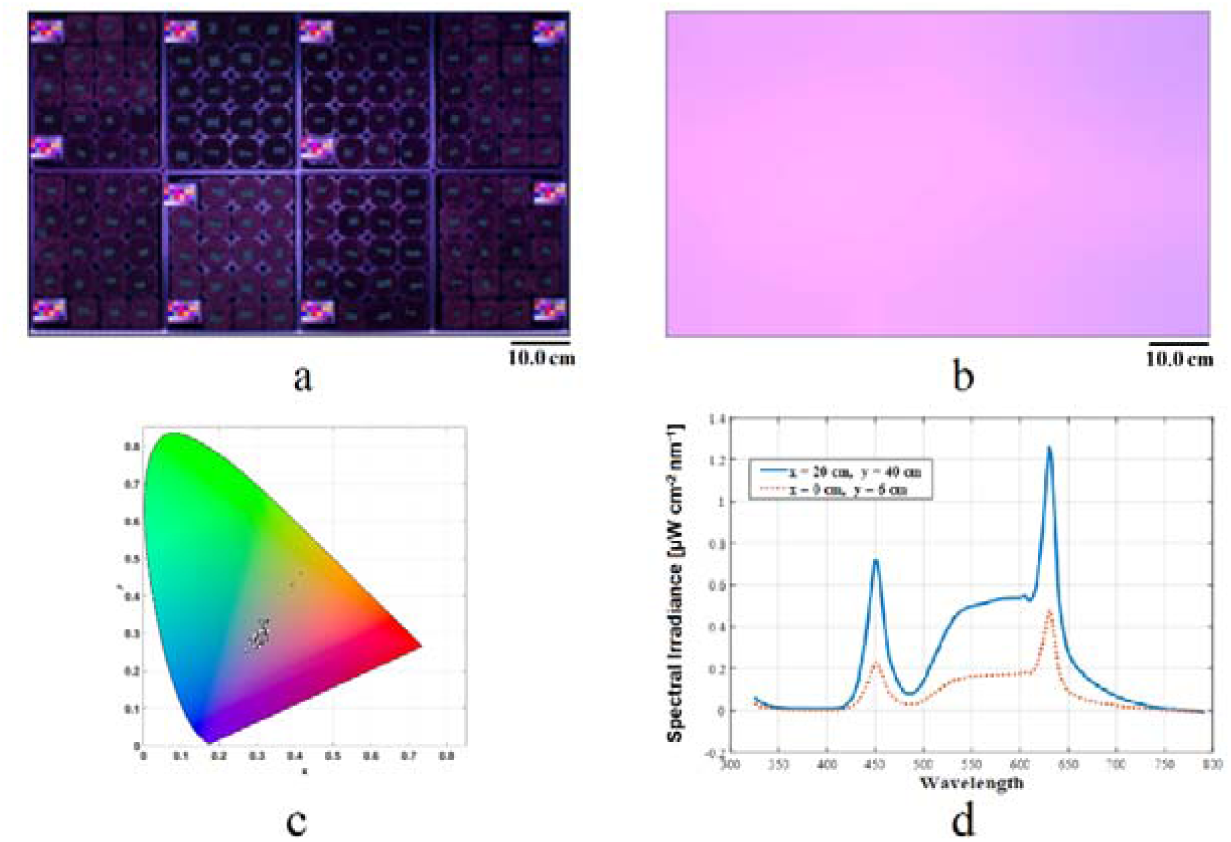
Light environment characteristics in the HTPP system. (a) A top-view image capturing one hundred and forty-eight pots and twelve Macbeth ColorCheckers, illustrating non-uniform illumination. (b) Spatial distribution of the illumination color on the plant phenotyping scene. (c) Illumination color distribution on the Chromaticity diagram using the 1931 CIE colorimetric system coordinates. (d) Spectral Irradiance at (x = 20 cm and y = 20 cm) and (x = 0 cm and y = 6 cm), the wavelength range is from 325 nm to 785 nm on the horizontal axis and the spectral irradiance [μW cm^-2^ nm^-1^] on the vertical axis. Two peaks represent the primary emission LED lights at blue (450 nm) and red (630 nm) and a curve plateau between these peaks (550 - 600 nm).

We created a ground truth image dataset to train and test our approach. This dataset consisted of 12 phenotyping images (5105 × 3075 pixels), which were illuminated with different light spectra and intensities using LED lights (**Figure 3**). A phenotyping image captured 148 pots and 12 Macbeth ColorCheckers, which were uniformly distributed on the image scene. Each pot contained a square of green fabric of known, uniform color. We extracted individual pot (290 × 290 pixel) and Macbeth ColorChecker (330 × 220 pixel) images by cropping them from phenotyping images. The resulting pot images showed a black plastic pot filled with soil and a green piece of fabric on its surface. The local ground-truth color was defined as the color of the fabric pieces as measured in the images. Therefore, this green fabric dataset provided the ground truth color on multiple parts of the picture, as the measured color varied across the scene.

**Figure 3.**
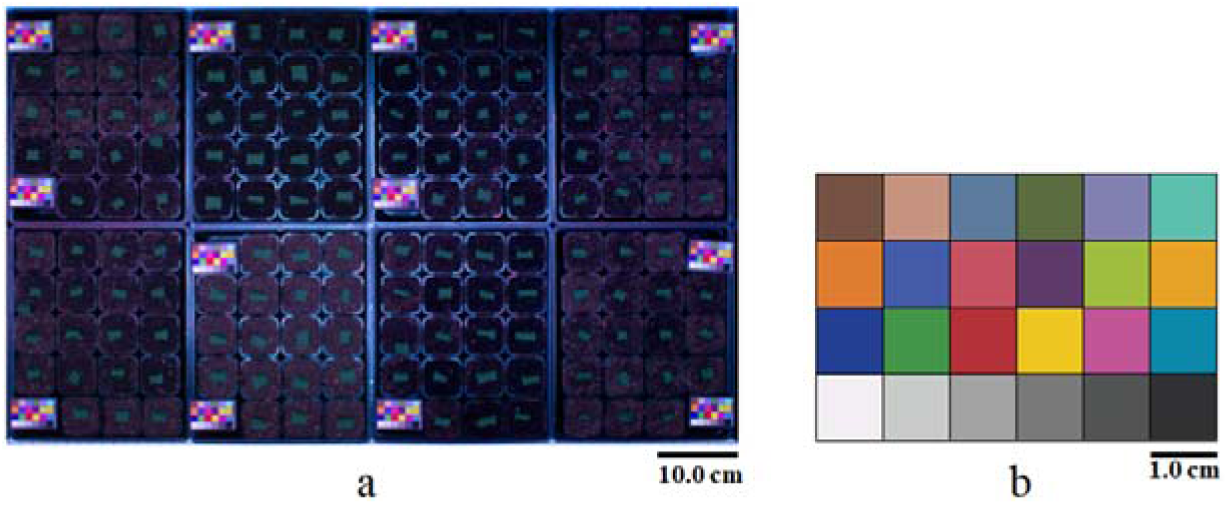
Green cloth ground truth dataset image sample. (a) The phenotyping image captured 148 pots and 12 Macbeth ColorCheckers, which were used to assess the algorithm performance. (b) Macbeth ColorChecker chart with 24 squares of defined reference color.

We applied our Bayesian color constancy algorithm on the green fabric dataset to correct the measured color of the green fabric back to its known value (**Figure 4**). We compared the performance of our algorithm (ColorBayes) with the calibration, grey world, white patch, and CLAHE algorithms (**Table 1**) [15-17]. To do so, we used the mean square error (MSE) to calculate the average of the square of the difference between ground-truth color values and each algorithm’s color estimation result. We calculated the MSE independently for each channel of the color image represented by the RGB color space. The Bayesian algorithm gave the most consistently low MSE values across the three image channels (R = 76.197, G = 110.647, B = 50.103), meaning it gave the highest performance in identifying the cloth’s true color. The calibration algorithm gave the next best performance (R = 66.443, G = 180.296, B = 244.659). The CLAHE algorithm had the highest MSE (R = 2809.441, G = 2531.786, B = 7674.055). The gray-world and white patch algorithms also had higher MSE values. A possible reason for high error among these latter three algorithms is that they correct images based only on the information contained in the pixel values without using the Macbeth Colorcheckers. In contrast, the Bayesian and calibration algorithms both estimate the illuminant properties by sampling the Macbeth ColorChecker. Accordingly, our results suggested that having a color reference within the image helped achieve higher color accuracy. Overall, the results indicate that our Bayesian algorithm improves color correction on a ground truth dataset relative to other algorithms commonly used in plant phenomics.

**Table 1.**
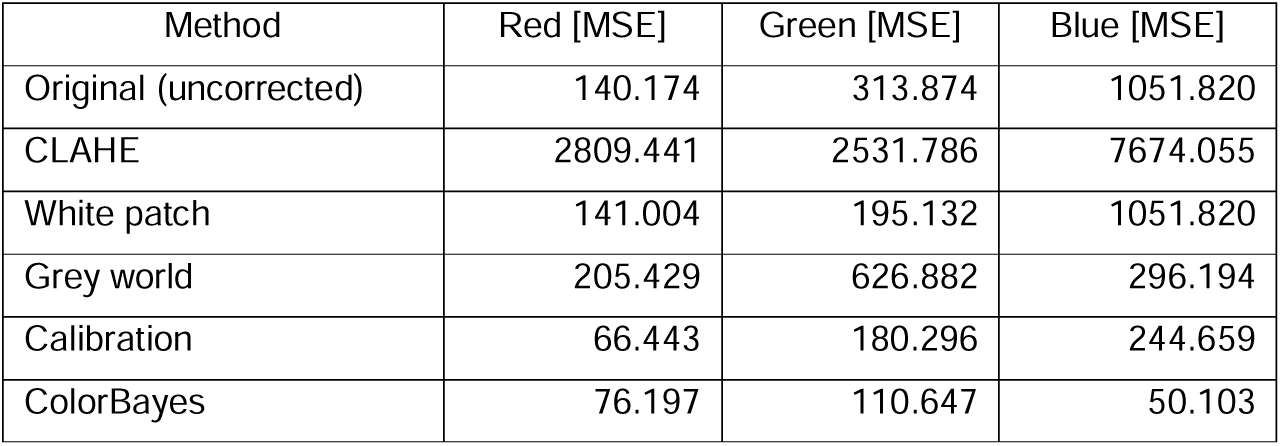
Accuracy color comparison between five color constancy algorithms on the green fabric ground-truth dataset. The numerical value is the mean square error representing the average squared difference between the estimated and ground-truth color values.

| Method | Red [MSE] | Green [MSE] | Blue [MSE] |
| --- | --- | --- | --- |
| Original (uncorrected) | 140.174 | 313.874 | 1051.820 |
| CLAHE | 2809.441 | 2531.786 | 7674.055 |
| White patch | 141.004 | 195.132 | 1051.820 |
| Grey world | 205.429 | 626.882 | 296.194 |
| Calibration | 66.443 | 180.296 | 244.659 |
| ColorBayes | 76.197 | 110.647 | 50.103 |

**Figure 4.**
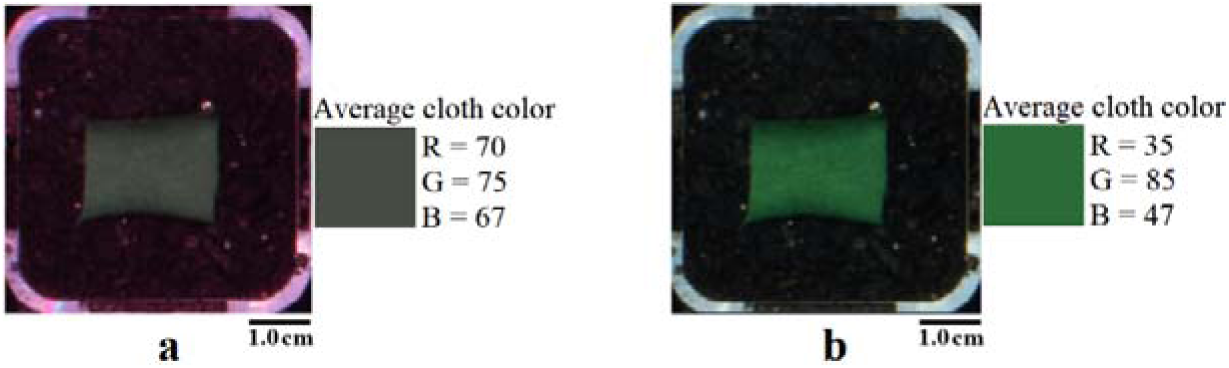
Color Bayes Bayesian color constancy results on the green cloth dataset. (a) shows the original pot image that was locally illuminated with a red cast light, (b) shows the same pot after applying our Bayesian algorithm.

Next, we applied the five algorithms to real plant datasets and assessed their performance using the MSE metric (**Table 2**). As with the green fabric ground truth dataset, our ColorBayes algorithm could effectively remove the local color cast from the pot images (**Figure 5**). It outperformed the other algorithms, obtaining the lowest MSE values in two image channels (R = 140.565, B = 143.50) and very close to the leading algorithm in the third channel (ColorBayes, G = 437.390; Calibration, 414.983). The calibration algorithm was the next-best performer (R = 296.207, G = 414.983, B = 572.333), followed by the Grey world, white patch, and CLAHE algorithms.

**Table 2.**
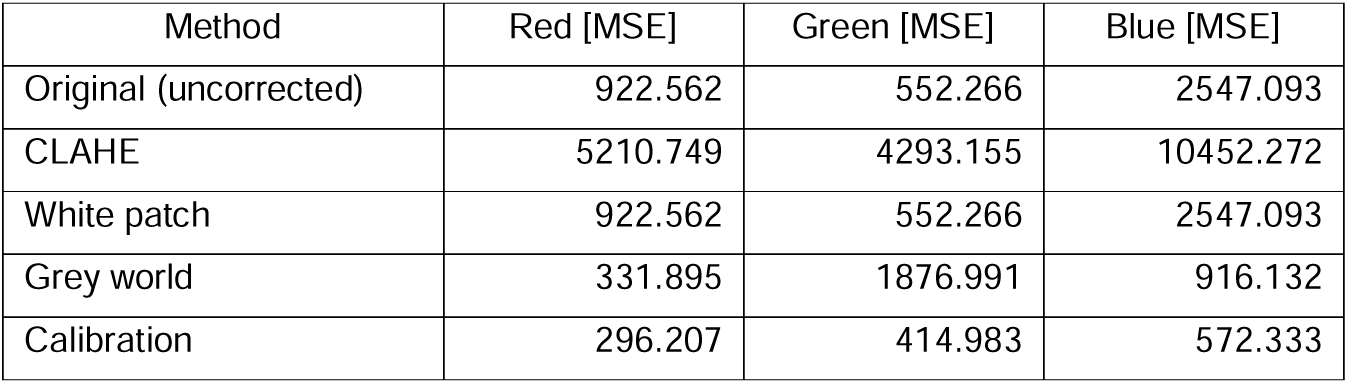
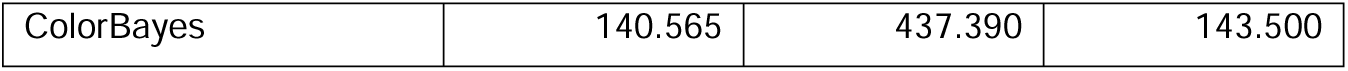
Accuracy of color comparison between five color constancy algorithms using real-world plant image data. The numerical value is the mean square error representing the average squared difference between the estimated and ground-truth color values.

**Figure 5.**
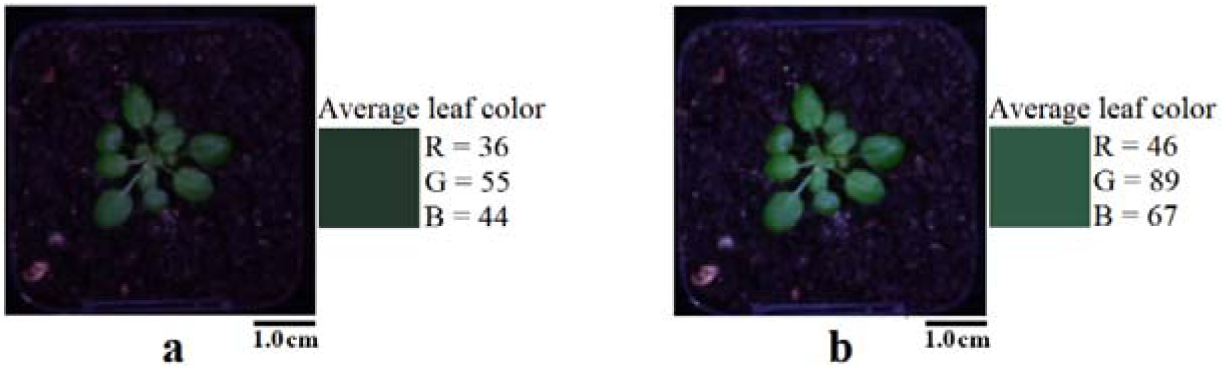
The Bayesian color constancy results on the real-world plant dataset. (a) shows the original pot image that was locally illuminated with low intensity and reddish illuminant light, (b) shows the same pot after applying our Bayesian algorithm.

In general, the MSE values were lower on the green fabric dataset than on the plant datasets, indicating better algorithm performance on the green fabric. This result is to be expected. Obtaining the true leaf color was not a trivial task as, unlike the green fabric, the plants in our experiment were not of identical color. In both cases, the dominant illuminant had comparable properties - light spectra and intensities that ranged from white light to saturated red, purple, and yellow light colors. However, the setup of the scene on the green cloth was the same in every image captured. This meant that the soil, cloth fabric, and pots had similar reflectance across the image dataset. On the other hand, the reflectance on the scenes in the plant dataset was more complex due to the natural color variation on leaves and the variation accumulated in the soil during cultivation. Each plant experiences a slightly different local environment that cannot be fully controlled, resulting in minor differences in plant development that are frequently reflected in slight variations in leaf color.

Detecting outlier plants within HTPP experiments can aid analysis, enabling individuals with interesting properties or suffering unintended disease to be identified [24]. The improvements in leaf color dispersion achieved by applying our algorithm enabled more sensitive detection of outliers (**Table 3**). Before color correction, the leaf color distribution was skewed, whereas the distribution was closer to normal after correction (**Figure 6**). We used the accuracy metric to evaluate the percentage of correct identification of outliers (calculated as the sum of true positives and true negatives, divided by all cases; **Table 3**). Fifteen true positives were identified by human-eye inspection for plants with abnormal color and development. The Bayesian-based algorithm had the highest accuracy (94.94%) as it homogenized the leaf color by removing the local color cast (**Figure 7**). Three out of fifteen outliers were detected based on plant color prior to application of the algorithm, indicating that false-negative results existed in the dataset. This occurred because prior to correction, the color of individual plants had higher divergence from the population plant color.

**Table 3.** Accuracy of outlier detection comparison between five color constancy algorithms using real-world plant image data.

| Method | True Positive | False Positive | True Negative | False Negative | Accuracy |
| --- | --- | --- | --- | --- | --- |
| Original (uncorrected) | 3 | 0 | 143 | 12 | 92.41% |
| Grey World | 3 | 0 | 143 | 12 | 92.41% |
| White patch | 3 | 0 | 143 | 12 | 92.41% |
| Calibration | 3 | 0 | 143 | 12 | 92.41% |
| CLAHE | 4 | 6 | 143 | 5 | 93.04% |
| Our Bayesian approach | 7 | 7 | 143 | 1 | 94.94% |

**Figure 6.**
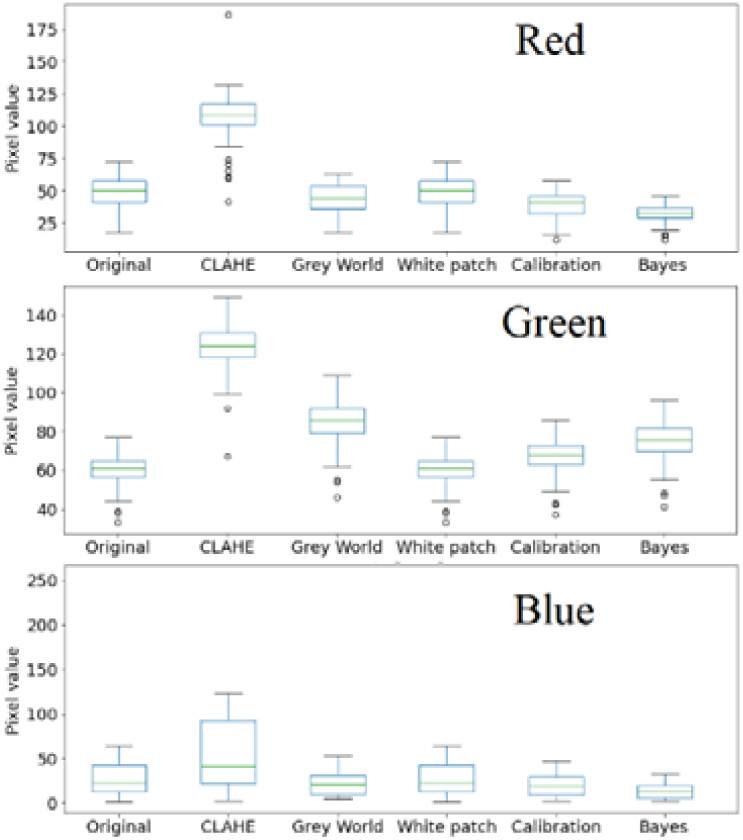
Comparing leaf color distribution after applying the five algorithms on the plant dataset at 29 DAS. The Bayesian algorithm reduced the variance among leaf color and identified more outliers than the other algorithms (CLAHE, grey world, white patch, and calibration). Original indicates non-color corrected dataset.

**Figure 7.**
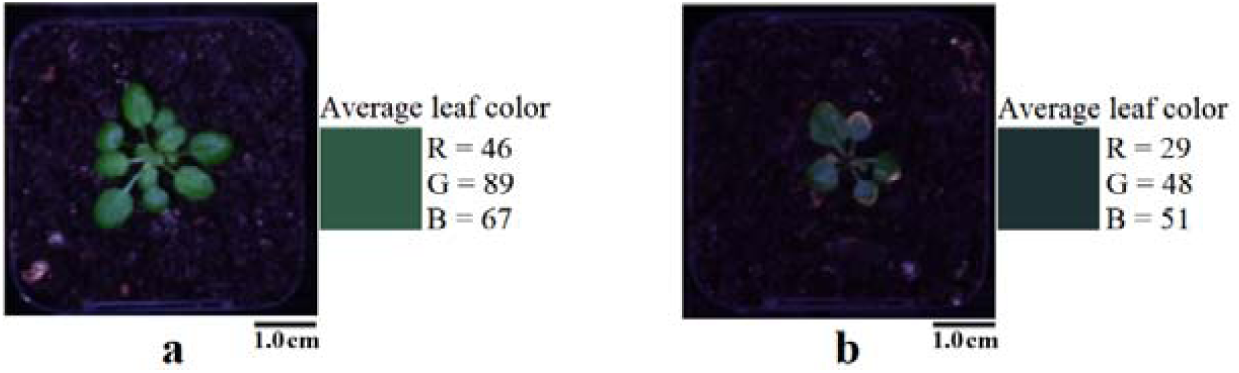
Color difference between a healthy and unhealthy plant. (a) healthy plant and its average color, (b) unhealthy plant and its average color.

## Discussion

Standard color constancy methods applied to image-based phenotyping rely on the assumption of a single and uniform illuminant. This assumption does not hold in typical HTPP systems because they have multiple, non-uniform illuminants such as LED arrays or fluorescent tubes. As a result, leaf colors may be incorrectly identified among plants within an experiment, which can mislead researchers and result in mistaken biological interpretations of data.

We propose a color constancy method, ColorBayes, based on Bayesian inference to correct the leaf color in plant phenotyping datasets. We tested this method using a ground truth color control dataset that contains pieces of green fabric and a manually annotated dataset of *A. thaliana* plants. Our method’s utility was demonstrated by correcting the leaf color of plants more accurately than commonly used color correction algorithms. The application of our algorithm will enable biologists to obtain more precise measures of plant color, permitting more reliable analyses of plant growth and development.

Our color constancy method improved outlier plant detection. Several factors can cause differences in leaf color between genetically identical plants grown in a single phenotyping experiment. These include small, uncontrollable local environmental effects that result in plant metabolic changes, stress and developmental responses, as well as image color distortion. If the image color distortion is reduced, it is possible to highlight leaf color changes produced by biological causes. However, multiple illuminants in large-scale experiments can locally cast the leaf color. Our method reduced the color distortion produced by technical issues and allowed the systematic detection of a greater number of outliers in the dataset than other, less accurate, color correction methods. We provide the code and instructions for applying ColorBayes freely at https://github.com/diloc/Color_correction.git.

## Methods

### Plant material and growth conditions

*A. thaliana* wild-type plants (ecotype Col-0) were grown in a chamber at 20□C with 50% humidity and watered every four days. The plants were kept in dark conditions for 10 hours and illuminated for 14 hours using LED lights (six colors) and cold white LED lights (∼4200□K) (**Table 4**). The light distribution on the growth chamber was not uniform in space and spectrum (**Figure 2**).

**Table 4.** LED light source specifications.

| LED light color | Wavelength range |
| --- | --- |
| Blue | 450-460 |
| Green | 530 |
| Red | 618-630 |
| Deep red | 660 |
| Far-red | 740 |
| Near-infrared | 850 |

### Image acquisition

The plants were imaged using top-view Digital Single-Lens Reflex (DSLR) cameras with a fixed location in the growing chamber operating in the visible light spectrum [24]. After acquisition, images were pre-processed to correct the perspective and lens distortion according to our previously published approach [24]. The image scene was divided segmented into individual pots. These areas had a similar size and similar objects such as plants, soil, pot container, and surroundings. Pixel colors varied between segmented images. The color difference in the plant dataset was driven by many factors, including the water content in the soil (wet-dry), the health and stage development of plants. Also, image colors varied from one location to another due to the non-uniform light distribution that locally cast it (**Figure 2**).

### Ground truth image dataset

An image dataset was created to estimate the light pattern in the HTPP system used to train and increase color accuracy (**Figure 3**). This image dataset was composed of RGB (300 × 300 pixel) images taken in the HTPP system. Each training image had similar elements as a common plant phenotyping image, including soil, pot, and surroundings, but a uniform green fabric replaced the plants. This replacement allowed minimal color variability among multiple images compared with natural plants as biological variations can influence leaf color. The training dataset assisted us in accounting for the color gradient produced by the system’s non-uniform lighting conditions. The algorithm learned the relationship between the local light conditions and color response on the image and subsequently estimated the actual color of the image under test.

### Image model

A digital image in the visible spectrum range ω can be represented by an array of pixels that contain the color information in three independent channels (red, green, blue). The pixel value in an image channel is defined in (**Eq. 1**). It depends on the spectral response of the camera Cλ at a particular wavelength λ, the energy of the light source E(λ), and the reflectance characteristics of the scene surfaces X(λ), which are assumed to be Lambertian surfaces.

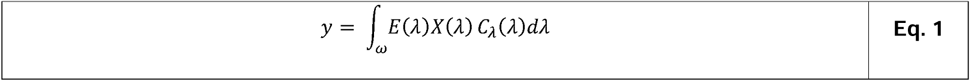

### Bayesian Color Constancy Approach

Our goal is to correct the local color distortion on plant phenotyping images caused by non-uniform illumination. The corrected image will show the colors of individual plants as if they were taken under the same standard illuminant (D65). This color constancy approach has two main steps. The first step is to estimate an unknown illuminant’s color and spatial distribution that causes the local color distortion. For this step, it is required a training dataset (ground truth), observed image data. Also, it is used the Bayes’ rule and the maximum a posteriori (MAP). The second step is to transform the observed image using the chromaticity adaptation method.

Before estimating the unknown illuminant, it is necessary to define the following assumptions:

1. An observed image (5105 × 3075 pixels) is made up of three independent color channels. (*k =* {*R, G, B*}) and divided into areas that correspond to individual pot areas.
2. A pixel class is assigned individually to segmented objects in a pot area such as plant and soil pixels. It means that a pixel class *Z*_*k*_ = {*z*_*ki*_} is a collection of pixels within the same spatial neighborhood and similar color values. The pixel value *z*_*ki*_ is a random variable at location *i =* 0,1,2,*…, n*.
3. The reflectance of a pixel class is a collection of reflectance *R*_*k*_ = {*r*_*ki*_}, Where *r*_*ki*_ is a random variable representing the reflectance at the location *i =* 0,1,2,*…, n*. Two adjacent reflectance are independent of each other, and the joint probability of is given by *p*(*r*_*ki*_, *r*_*kj*_) = *p*(*r*_*ki*_)*p*(*r*_*kj*_). Based on the same assumption, all reflectance in a pixel class are independent events with joint probability 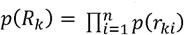.
4. The illuminant of a pixel class is a collection of illuminants *L*_*k*_ = {*l*_*ki*_}. However, it is assumed that the illuminant is constant for all pixels in a class, meaning, *l*_*k*_ = *l*_*ki*_ and *L*_*k*_ = {*l*_*k*_}. Then, the probability distribution of the illuminant is uniform, *p*(*l*_*k*_) = *u*_*k*_, being *u*_*k*_ a constant value.
5. The illumination and the reflectance are statistically independent of each other *p*(*L*_*k*_, *R*_*k*_) = *p*(*L*_*k*_)*p*(*R*_*k*_).
6. The value of a pixel *Z*_*ki*_ is a function of the reflectance *r*_*ki*_, the illuminant *l*_*ki*_ and the Gaussian noise *w*_*ki*_ with a mean equal to zero and variance 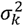 (**Eq. 2**).

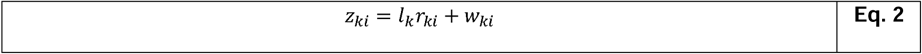

The multivariable function described in **Eq. 2** can be statistically represented using the likelihood function. It is equivalent to Gaussian noise probability distribution (**Eq. 3**).

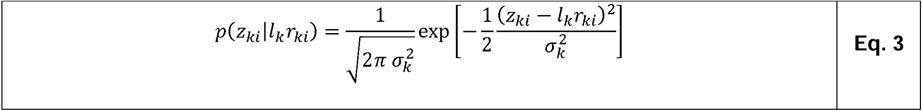

### Priors

The reflectance prior distribution is informative and obtained from an existing training dataset. It assembles a normal probability distribution with mean equal to *μ*_*k*_ and variance 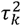 (**Eq. 4**).

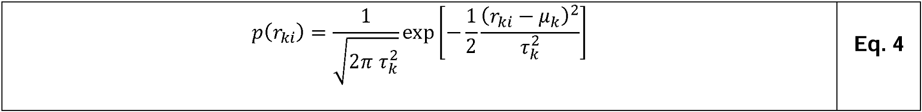

The illuminant prior distribution is also obtained from the training data and follows a uniform distribution over a pixel class, where μ_*k*_is constant (**Eq. 5**).

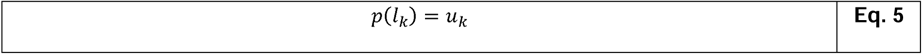

### Posterior

The posterior is based on the Bayes’ theorem, and it depends on two variables in our case (*l*_*k*_,*r*_*ki*_). It is possible to simplify the problem by marginalizing the posterior concerning the reflectance variable (**Eq. 6**). The reflectance is considered a nuisance variable with no particular interest in being estimated.

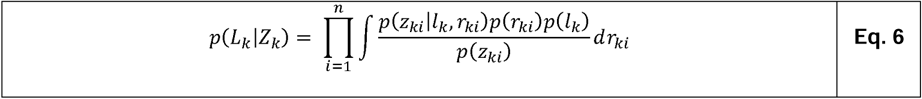

The next step is to replace terms in **Eq. 6** by **Eq. 3, Eq. 4**, and **Eq. 5**, as shown in **Eq. 7**.

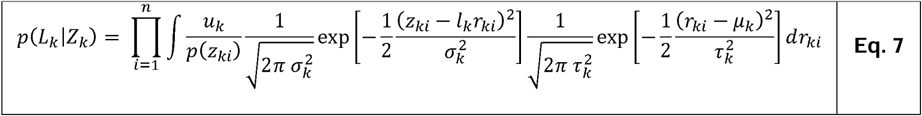

The posterior is solved by simplifying the exponent, expanding the product function, and using the Euler–Poisson integral solution (**Eq. 8**).

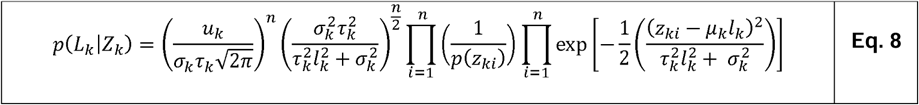

The next step is to find an illuminant *l*_*k*_ where the fixed-shape distribution in **Eq. 8** reaches the maximum value. However,there is no interest in estimating the maximum value of the distribution itself. Under this assumption, the first three terms of **Eq. 8** can be treated as a scale factor for the exponential function. It means that the posterior distribution can be reduced to a proportional relationship (**Eq. 9**).

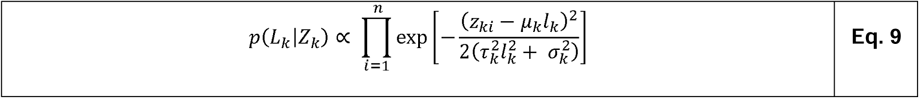

Next step is to apply the natural logarithm on the two sides of the proportional relationship (**Eq. 10**).

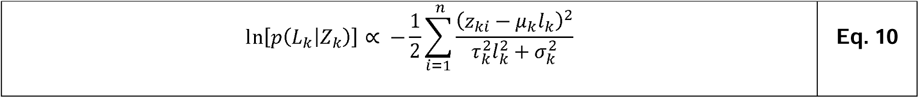

The following step is to calculate the critical point by applying the first derivate concerning the illuminant variable (*l*_*k*_) and make it equal to zero (**Eq. 11**). The critical point gives the *l*_*k*_ value where the exponential function reaches it maximum value.

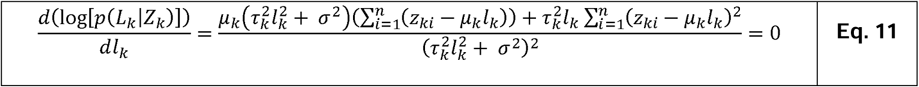

The denominator of the previous rational expression is a positive real number if the pixel class variance, prior variance, and the illuminant value are greater than zero. This condition is met in our approach. So, if the rational expression is equal to zero, then the numerator must be zero. It means that the Maximum a posteriori is given by **Eq. 12**.

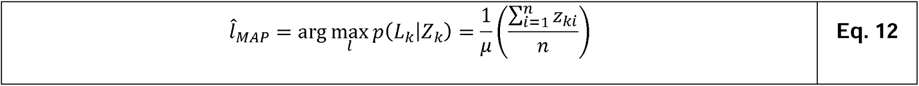

### Illuminant surface estimation

A local illuminant gives only an estimation for a limited portion of an image where the pixel class is located. It is necessary to calculate an illuminant surface that covers the whole image. The illuminant surface recreates the spatial illumination pattern on the scene. Two linear spatial interpolators were used to estimate the illuminant values between pixel classes. A two-kernel Gaussian filter is applied to the illumination surface to adjust the intensity and preserve the illumination soft spatial transition.

### Chromaticity adaptation

After calculating the spatial illumination distribution for an image, we use the chromaticity adaptation method to correct the image colors (**Eq. 13**). It reduces the local color cast produced by different illumination conditions within an image. Also, it moves the image colors closer as the image was taken under a spatial uniform D65 standard illuminant.

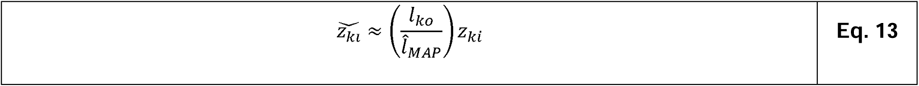

## Abbreviations

CER: Controlled environment room
CLAHE: Contrast limited adaptive histogram equalization
DAPD: Digital adjustment of plant development
DAS: Day-after-sowing
DSLR: Digital single-lens reflex
HSV: Hue, saturation, value
HTPP: High throughput plant phenotyping
LED: Light-emitting diode
PAR: Photosynthetically active radiation
RGB: Red, green, blue

## Authors’ contributions

Experiment design: DL, JW, and MGL. DL performed the experiments. Data acquisition and analysis: DL, GD, EC, and MGL. Interpreted the results: DL and MGL. All authors read and approved the final manuscript.

## Acknowledgments

We thank Dr. Ricarda Jost and Dr. Oliver Berkowitz for their valuable contribution to the plant experiment design.

## Competing interests

The authors declare that they have no competing interests.

## Availability of data and materials

The code of the color correction algorithm is available for reuse at https://github.com/diloc/Color_correction.git. Correspondence should be addressed to.

## Consent for publication

All authors reviewed and approved the final version of the manuscript for submission.

## Ethics approval and consent to participate

Not applicable.

## Funding

Work in the Lewsey and Whelan labs was funded by the Australian Research Council Industrial Transformation Hub in Medicinal Agriculture (IH180100006). Work in the Whelan Lab was funded by the Australian Research Council Centre of Excellence in Plant Energy Biology (CE140100008). DL was the recipient of scholarships from La Trobe University Graduate Research School.

## Notes

https://github.com/diloc/Color_correction.git

### Competing Interest Statement

The authors have declared no competing interest.

## References

1. Casadesús J, Kaya Y, Bort J, Nachit MM, Araus JL, Amor S, Ferrazzano G, Maalouf F, Maccaferri M, Martos V et al: Using vegetation indices derived from conventional digital cameras as selection criteria for wheat breeding in water-limited environments. Annals of Applied Biology 2007, 150(2):227–236.

2. Serret MD, Al-Dakheel AJ, Yousfi S, Fernáandez-Gallego JA, Elouafi IA, Araus JL: Vegetation indices derived from digital images and stable carbon and nitrogen isotope signatures as indicators of date palm performance under salinity. Agricultural Water Management 2020, 230:105949.

3. Menzies IJ, Youard LW, Lord JM, Carpenter KL, van Klink JW, Perry NB, Schaefer HM, Gould KS: Leaf colour polymorphisms: a balance between plant defence and photosynthesis. Journal of Ecology 2016, 104(1):104–113.

4. Sims DA, Gamon JA: Relationships between leaf pigment content and spectral reflectance across a wide range of species, leaf structures and developmental stages. Remote Sensing of Environment 2002, 81(2-3):337–354.

5. Uchida R: Essential nutrients for plant growth: nutrient functions and deficiency symptoms. Plant nutrient management in Hawaii’s soils 2000:31–55.

6. Tucker AO, Maciarello MJ, Tucker SS: A Survey of Color Charts for Biological Descriptions. Taxon 1991, 40(2):201–214.

7. Hunt ER, Cavigelli M, Daughtry CST, McMurtrey JE, Walthall CL: Evaluation of Digital Photography from Model Aircraft for Remote Sensing of Crop Biomass and Nitrogen Status. Precision Agriculture 2005, 6(4):359–378.

8. Chopin J, Kumar P, Miklavcic SJ: Land-based crop phenotyping by image analysis: consistent canopy characterization from inconsistent field illumination. Plant Methods 2018, 14(1):39.

9. Sadeghi-Tehran P, Virlet N, Sabermanesh K, Hawkesford MJ: Multi-feature machine learning model for automatic segmentation of green fractional vegetation cover for high-throughput field phenotyping. Plant Methods 2017, 13(1):103.

10. Casadesús J, Biel C, Savé R: Turf color measurement with conventional digital cameras. In: EFITA/WCCA Joint Congress in Agriculture: 2005. Universidade de Trás-os-Montes e Alto Douro Vila Real: 804–811.

11. Luo MR: Encyclopedia of color science and technology: Springer Reference; 2016.

12. Foster DH: Color constancy. Vision Research 2011, 51(7):674–700.

13. Vazquez-Corral J, Párraga C, Baldrich R, Vanrell M: Color constancy algorithms: Psychophysical evaluation on a new dataset. Journal of Imaging Science and Technology 2009, 53(3):31105-31101- 31105-31109.

14. Gijsenij A, Lu R, Gevers T: Color Constancy for Multiple Light Sources. IEEE Transactions on Image Processing 2012, 21(2):697–707.

15. Buchsbaum G: A spatial processor model for object colour perception. Journal of the Franklin Institute 1980, 310(1):1–26.

16. Gevers T, Gijsenij A, van de Weijer J, Geusebroek J-M, Geusebroek J-M: Color in Computer Vision : Fundamentals and Applications. Somerset, UNITED STATES: John Wiley & Sons, Incorporated; 2012.

17. Tang S, Dong M, Ma J, Zhou Z, Li C: Color image enhancement based on retinex theory with guided filter. In: 2017 29th Chinese Control And Decision Conference (CCDC): 28-30 May 2017 2017. 5676–5680.

18. Brainard DH, Freeman WT: Bayesian color constancy. Journal of the Optical Society of America A, Optics, image science, and vision 1997, 14(7):1393–1411.

19. Rosenberg C, Minka T, Ladsariya A: Bayesian color constancy with non-Gaussian models. In: Advances in Neural Information Processing Systems: 2004.

20. Gehler PV, Rother C, Blake A, Minka T, Sharp T: Bayesian color constancy revisited. In: 2008 IEEE Conference on Computer Vision and Pattern Recognition: 23-28 June 2008 2008. 1–8.

21. Wransky M: True color measurements using color calibration techniques. In. edited by Espanol M, Wilber JP, Ye J: ProQuest Dissertations Publishing; 2015.

22. Ojo JA, Solomon ID, Adeniran SA: Contrast enhancement algorithm for colour images. In: 2015 Science and Information Conference (SAI): 28-30 July 2015 2015. 555–559.

23. Martinelli T, Potenza E, Moschella A, Zaccheria F, Benedettelli S, Andrzejewska J: Phenotypic Evaluation of a Milk Thistle Germplasm Collection: Fruit Morphology and Chemical Composition. Crop Science 2016, 56(6):3160–3172.

24. Lozano-Claros D, Meng X, Custovic E, Deng G, Berkowitz O, Whelan J, Lewsey MG: Developmental normalization of phenomics data generated by high throughput plant phenotyping systems. Plant Methods 2020, 16(1):111.

